# Systematic identification of significantly mutated regions reveals a rich landscape of functional molecular alterations across cancer genomes

**DOI:** 10.1101/020875

**Authors:** Carlos L. Araya, Can Cenik, Jason A. Reuter, Gert Kiss, Vijay S. Pande, Michael P. Snyder, William J. Greenleaf

**Author notes:** These authors contributed equally to this work (Carlos L. Araya, Can Cenik).

## Abstract

**One Sentence Summary:** Identification of multi-scale mutational hotspots in cancer exomes facilitates understanding of mutations both within coding and non-coding elements.

Cancer genome sequencing studies have identified cancer-driver genes from the increased accumulation of protein-altering mutations. However, the positional distributions of coding mutations, and the 79% of somatic variants in exome data that do not alter protein sequence or RNA splicing, remain largely unstudied. We employed density-based clustering methods on ∼4,700 exomes from 21 tumor types to detect variably-sized significantly mutated regions (SMRs). SMRs reveal recurrent alterations across a diverse spectrum of coding and non-coding elements, including microRNAs, transcription factor binding sites, and untranslated regions that are individually mutated in up to ∼15% of samples in specific cancer types. SMRs often associated with changes in gene expression and signalling. Mapping SMRs to protein structures revealed spatial clustering of somatic mutations at known and novel cancer-driver domains and molecular interfaces. Mutation frequencies in SMRs demonstrate that distinct protein regions are differentially mutated among tumor types. The functional diversity of SMRs underscores both the varied mechanisms of oncogenic misregulation and the advantage of unbiased driver identification.

---

In cancer, somatic driver mutations alter functional elements of diverse nature and size. For example, melanoma drivers include hyper-activating mutations at single amino acid residues (e.g. BRAF V600^1^), inactivating mutations along tumor suppressor exons (e.g. *PTEN*^1^), and regulatory mutations (e.g. *TERT* promoter^2^). Cancer genomics projects, such as the The Cancer Genome Atlas (TCGA) and the International Cancer Genome Consortium (ICGC), have substantially expanded our understanding of the landscape of somatic alterations by identifying frequently mutated protein coding genes^3–5^. However, these studies have focused little attention on systematically analyzing the positional distribution of coding mutations or characterizing non-coding alterations^6^.

Most algorithms to identify cancer-driver protein-coding genes examine non-synonymous to synonymous mutation rates across the gene body or recurrently mutated amino acids called mutation hotspots^5^, as observed in BRAF^7^, IDH1^8^, and DNA polymerase ε (POLE)^9^. Yet, these analyses ignore recurrent alterations in the vast intermediate scale of functional coding elements, such as protein subunits or interfaces. Moreover, where mutation clustering within genes has been examined^10–12^, analyses have employed fixed base-pair windows or identified clusters of non-synonymous mutations, assuming driver mutations exclusively impact protein sequence and ignoring the importance of exon-embedded regulatory elements^13–17^.

Indeed, a significant proportion of regulatory elements in the genome occurs in, or proximal to, exons^14,18^, suggesting many may be captured by whole-exome sequencing (WES). Such data makes the investigation of regulatory elements especially attractive, as our understanding of non-coding mutations in cancer remains significantly underdeveloped, despite clear examples of importance (i.e. *TERT* promoter). Recent efforts to begin to characterize non-coding variation in cancer genomes have examined either (1) pan-cancer whole-genome sequencing (WGS) data, or (2) predefined regions –such as ETS binding sites, splicing signals, promoters, and untranslated regions (UTRs)– or mutation types^19–21^. These approaches either presume the relevant targets of disruption, or disregard the established heterogeneity among tumor types at the level of cancer-driver genes and pathways^5,22^, as well as in nucleotide-specific mutation probabilities^3,4^. Systematic analyses of metazoan regulatory activity have revealed substantial tissue and developmental stage specificity^23–25^, suggesting that mutations in cancer-type-specific regulatory features may be significant non-coding drivers of cancer.

To address these diverse limitations, we employed density-based spatial clustering techniques utilizing cancer and gene-specific mutation models to identify and assess the significance of clusters of recurrent coding and non-coding mutations in 21 cancer types. This approach permitted the unbiased identification of variably-sized genomic regions recurrently altered by somatic mutations, which we term significantly mutated regions (SMRs). We identified SMRs in numerous well-established cancer-drivers as well as in novel genes and functional elements. Moreover, SMRs were associated with non-coding elements, protein structures, molecular interfaces, and transcriptional and signaling profiles, providing insight into the molecular importance of accumulating somatic mutations in these regions. Overall, SMRs revealed a rich spectrum of coding and non-coding elements recurrently targeted by somatic alterations that complement gene-centric analyses.

## Results

### Multi-scale detection of regions of high mutation density in cancer exomes

We re-annotated ∼3 million previously identified^5^ somatic, single nucleotide variants (SNVs) from 4,735 cancers of 21 tumor types (**Supplementary Fig. 1a**). We recorded^26^ the impact of each mutation on protein-coding sequences, other transcribed sequences, and adjacent regulatory regions (**Supplementary Fig. 1b**). Fully 79.0% (*n*=2,431,360) of these somatic mutations do not alter protein-coding sequences or their splicing and thus were not previously considered in the analysis of cancer-driver mutations^5^ (**Fig. 1a**).

**Figure 1.**
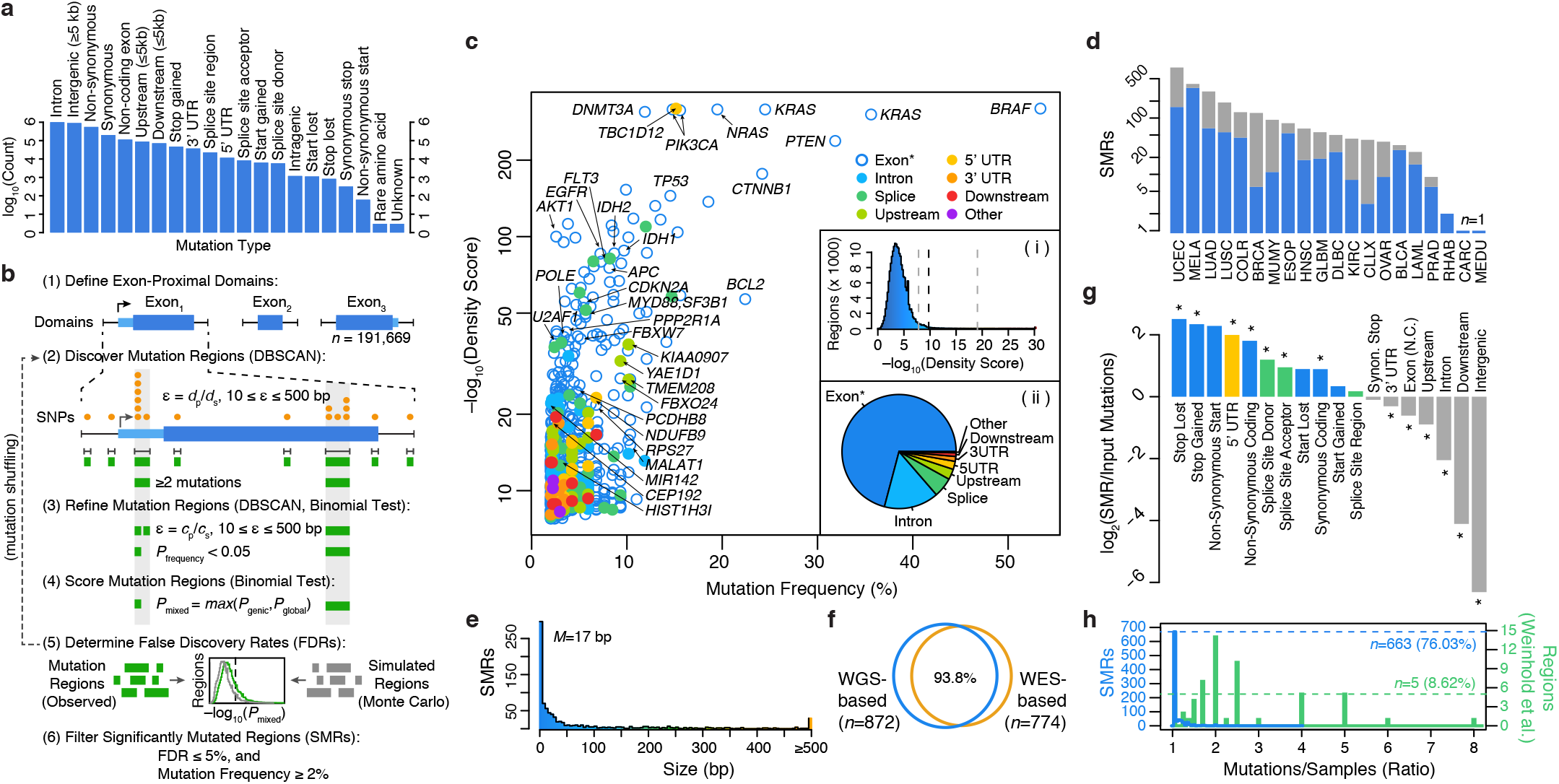
Identification of significantly mutated regions (SMRs) in 20 cancer-types across a broad spectrum of functional elements. (**a**) Pan-cancer distribution of mutation types in *n*=3,078,482 somatic single-nucleotide variant (SNV) calls. (**b**) SMR identification workflow. Exons and exon proximal domains (≤1,000 bp; blue) were scanned for clusters of somatic mutations (orange) with DBSCAN. Distance parameter ε is dynamically defined as the average distance of mutated positions (*d*_p_) in the domain size (*d*_s_), constrained within 10 ≤ ε ≤ 500 bp. Identified clusters (green) are divided if sub-clusters with higher (*P* < 0.05, binomial test) mutation densities are found in a second-pass DBSCAN analysis with ε defined as the average distance of mutated positions (*c*_p_) within the cluster of size *c*^s^. Cluster density scores are computed using the more conservative of the gene-specific and global background mutation rates as the combined binomial probabilities of the observed mutation density. For each cancer type, density score FDRs are computed by randomizing mutation positions (Supplementary Methods). SMRs were identified as clusters with FDR ≤5% density scores and mutated in ≥2% of cancer-specific samples. (**c**) Density scores and mutation frequencies of *n*=872 SMRs in 20 cancer types. SMRs are color-coded by region type. The distribution of density scores in evaluated regions and SMR region types are shown in insets (i) and (ii), respectively. Dashed lines indicate the minimum, median, and maximum density score FDR (5%) thresholds. (*) Exon label indicates coding regions and non-coding genes. (**d**) Number of SMRs with *FDR* ≤ 5% and mutation frequency ≥2% per cancer-type. Gray bars indicate the number of regions with *FDR* ≤ 5%, detailing the effect of the mutation frequency threshold. (**e**) SMR size distribution (median = 17 bp). (**f**) Concordance between SMRs discovered employing whole-genome sequence (WGS)-based and whole-exome sequence (WES)-based background models. (**g**) Fold change in mutation type representation between SMR-associated and input mutations. Asterisks denote categories with significant changes in representation (*P* < 0.01). Enriched mutation type colors match region types in (**c**). (**h**) Distribution of the mutations contributed per sample in SMRs (blue) and 58 (green) recurrently-altered non-coding regions^19^.

To systematically discover both coding and non-coding cancer-drivers, we applied an annotation-independent, density-based clustering technique^27^ to identify 198,247 variably-sized clusters of somatic mutations within exon-proximal domains of the human genome (**Fig. 1b**; Supplementary Methods). Notably, we included synonymous mutations within coding regions because functionally important non-coding features such as miRNAs^13^, regulatory RNA features^28^, and transcription factor (TF) binding sites^14^ can be embedded within these regions.

Mutation density scores within each identified cluster were derived as the Fisher’s combined *p*-value of the individual binomial probabilities of observing *k* or more mutations for each mutation type within the region across independent samples of each cancer type (Supplementary Methods). We evaluated mutation density for each cluster using gene-specific and genome-wide models of mutation probability (**Supplementary Fig. 2**), which were well-correlated (**Supplementary Fig. 3a**), and selected the more conservative estimate for each cluster as the final density score (Supplementary Methods). Gene-specific mutation probability models accounted for sequence composition (GC-content) as well as differences in local gene expression and replication timing, which have been shown to correlate with somatic mutation rate^4^. To avoid skewed mutation probability estimates due to selection pressure on exons, we applied a Bayesian framework to derive gene-specific mutation probabilities given intronic mutation probabilities in cancer WGS data^3,19^ while controlling for differences in sensitivity in WES and WGS (Supplementary Methods).

Increasing density scores correlated with stronger enrichments (up to 120x) for somatic SNV-driven cancer genes (*n*=158) as determined by the Cancer Gene Census (CGC)(**Supplementary Fig. 3b**)^29,30^. Although most somatic SNV-driven cancer genes do not display signals of high somatic mutation density (**Supplementary Fig. 3c**), ∼10% of genes associated with regions of extreme density scores (*P* ≤ 10^−20^) were not found previously in a gene-level analysis^5^ or in the CGC. Thus, high density scores are enriched for known cancer genes but also nominate novel cancer-driver genes.

We applied Monte Carlo simulations to select density score thresholds that control the false discovery rate (FDR) to ≤ 5% (**Supplementary Fig. 4, Supplementary Table 1**). We selected 872 regions (**Fig. 1c**), termed *Significantly Mutated Regions* (SMRs), that were altered in ≥2% of patients in 20 cancer types for further characterization (**Fig. 1d, Supplementary Fig. 5**). SMRs span 735 genomic regions, which are assigned unique SMR codes (e.g. *TP53.1*); note that some SMRs appear in more than one cancer type. We classified SMRs into “high”, “medium”, and “low” confidence sets on the basis of their density scores and contribution from mutator samples (**Supplementary Table 2**, Supplementary Methods). We observed correspondingly high (63.3x, *P* = 2.5 × 10^−46^), medium (6.2x, *P* = 2.6 × 10^−10^), and low (5.0x, *P* = 5.0 × 10^−4^) enrichments for somatic SNV-driven cancer genes in these sets. Over 87% of SMRs were contained within mappable (100 bp) regions of the genome, and an analysis of 6,179 recently-published breakpoints from 7 cancer types^31^ yielded a single SMR (in PTEN) within 50 bp of a resolved breakpoint, suggesting that the observed mutation density in SMRs is not attributable to mapping artifacts.

SMRs displayed a wide range of sizes (**Fig. 1e**, *median* = 17 bp), are robust to distinct mutation background models (**Fig. 1f**, Supplementary Methods), and are enriched in protein-coding, 5’ UTR and splice-site mutations (**Fig. 1g**, *P* < 0.01). Importantly, SMRs are not driven by samples that contribute large numbers of mutations per region (**Fig. 1h**). This is in contrast to recently proposed regions of recurrent alteration^19^ where as little as five were driven exclusively by distinct tumor samples (*P* = 6.0 × 10^−45^, Wilcoxon rank sum test). Thus, we have identified a functionally diverse set of variably-sized SMRs targeted by recurrent somatic alterations.

### SMRs are enriched in known cancer-drivers and implicate many novel cancer genes

SMRs, which harbor a diverse representation of predicted functional impacts on 610 genes, are significantly enriched in known cancer-driver genes (Lawrence et al.^5^ or CGC, *P* = 1.3 × 10^−34^, hypergeometric test), affecting a total of 91 known cancer-driver genes, including canonical oncogenes (e.g. *BRAF*, *KRAS*, *NRAS*, *PIK3CA*, and *CTNNB1*) and tumor suppressors (e.g. *PTEN*, *TP53*, and *APC*). SMR-associated genes also include 17 CGC genes previously undetected in a gene-level analysis^5^, such as established oncogenes like *BCL2* and *PIM1* and the cancer-associated non-coding gene *MALAT1*. Most coding region SMRs are driven by protein altering mutations (**Supplementary Fig. 6**), demonstrating coding SMRs capture positive selection primarily acting on protein alterations. In total, SMRs implicate 26 known cancer-driver genes to an additional 31 gene-to-cancer type associations not uncovered by a gene-level analysis^5^ (**Supplementary Table 3**). We note, however, that most known cancer-driver genes do not harbor regions of dense mutation recurrence within this data (see Supplementary Discussion, **Supplementary Fig. 7**), suggesting that SMR identification complements gene-level approaches.

We discovered SMRs in multiple novel cancer-driver genes, including the breast cancer-associated antigen and putative transcription factor ANKRD30A^32^, in which ∼21% of melanomas harbor mutations within one or more of three SMRs. Mutations in these SMRs were validated in WGS data from 6 of 17 cutaneous melanomas^3,19^. Within the entire gene-body, 27 of 118 WES and 10 of 17 WGS datasets from melanoma patients harbor somatic protein-altering mutations in ANKRD30A. Overall, of the 185 high confidence SMRs, 16 were associated with novel cancer-driver genes (**Supplementary Table 4**). As expected on the basis of methodological differences, these putative novel cancer-drivers are primarily (∼81%) driven by non-coding alterations, as discussed in the next section.

### SMRs implicate diverse non-coding regulatory features

A significant proportion (31.2%; *P* < 2.2 × 10^−16^, proportions test) of SMRs are not predicted to affect protein sequences, highlighting the potential for the discovery of pathological non-coding variation in WES data. In total, 130 SMRs lay within DNase I hypersensitive (DHS) sites^25^ and are enriched in promoter (*Q* = 4.0 × 10^−9^) and 5’ UTR features (*Q* = 4.4 × 10^−10^; **Supplementary Table 5**). Three promoter SMRs (*n*=29) coincide with regions deemed significantly mutated in a pan-cancer analysis of WGS data^19^. Across all cancer types, small (≤25 bp) non-coding SMRs were enriched in binding sequences for ETS oncogene family (*Q* = 2.6 × 10^−6^) and winged-helix repressor (*Q* = 2.0 × 10^−4^) TFs (**Fig. 2a**, **Supplementary Table 6**). We also detected cancer-specific TF motif enrichments within SMRs from diffuse large B-cell lymphoma, melanoma, and rhabdosarcoma (**Fig. 2b**, **Supplementary Table 7**).

**Figure 2.**
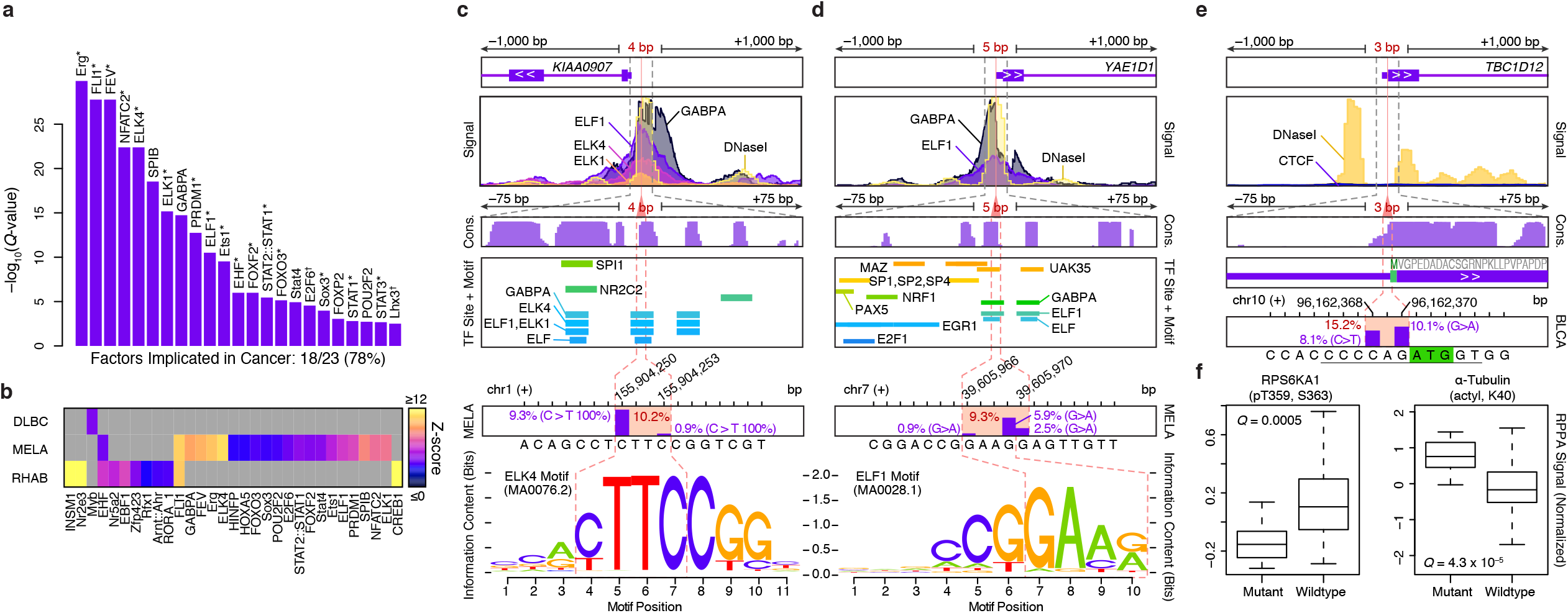
Non-coding SMRs recurrently alter promoters and 5’ UTRs. (**a**) Transcription factors (TFs) with enriched (*Q* < 0.01) motifs in small SMRs (≤25bp) across all cancer types are shown. 18 of the 23 TFs are known cancer-associated TFs (*) or associated with cell-cycle control or developmental roles (†). (**b**) Cancer-specific motif enrichment analysis. Gene structure, ENCODE ChIP-seq and DNaseI signals, vertebrate conservation (phastCons 100way), Factorbook TF binding sites and motif occurrences, and somatic mutation frequencies at melanoma SMRs in *KIAA0907* (**c**) and *YAE1D1* (**d**) promoter regions are shown at multiple scales (±1,000, ±75, and ±7 bp). Fraction of melanoma samples altered (mutation frequency) within each SMR (red) and at each position (purple bars) are shown. Motifs of *in vivo* ETS-family binding sites that overlap the SMRs are highlighted. (**e**) Gene-structure, ENCODE CTCF and DNaseI signals, vertebrate conservation (phastCons 100way), and protein coding sequence at the 5’ UTR *TBC1D12* bladder cancer SMR are shown at multiple scales. Start codon position is highlighted in green and Kozak sequence is underlined. CTCF signal is shown on the basis of a Factorbook CTCF site overlapping this SMR. (**f**) Relative protein and post-translational modification signals of wildtype and mutant (*TBC1D12.1* SMR-altered) bladder tumors.

We discovered (4 and 5 bp) SMRs within DHS sites of the *KIAA0907* and *YAE1D1* promoters that were altered in 10.2% and 9.3% of WES melanomas (**Fig. 2c,d**), respectively. Somatic mutations in these SMRs were confirmed in WGS data of melanomas (*n*=1 for *KIAA0907* and *n*=2 for *YAE1D1* of *n*=17, respectively)^3,19^. Yet, these regions did not reach significance in a pan-cancer analysis^19^, highlighting cancer-specificity in non-coding alterations. In both SMRs, mutations alter core-recognition sequences within *in vivo* ETS factor binding sites, with varying effects on ETS primary sequence preferences. *KIAA0907* encodes a largely uncharacterized putative RNA-binding protein. However, intronic sequences in this gene harbor *SNORA42*, an H/ACA class snoRNA with increased expression in lung cancer^33^, suggesting promoter SMR alterations may enhance transcription at this locus. RNA-level overexpression of *YAE1D1* has previously been observed in lower crypt-like colorectal cancer^34^, and a small cohort of melanoma samples showed increased YAE1D1 protein levels compared to untransformed melanocytes^35^, suggesting that *YAE1D1* may also be upregulated in melanomas.

In addition to SMRs that impact promoter regions, we observed 32 SMRs in 5’ and 3’ UTRs. Most strikingly, we discovered a 3 bp SMR in the 5’ UTR of *TBC1D12* that is mutated in ∼15% of bladder cancers (**Fig. 2e**). Recurrent mutations were positioned near the start codon (Kozak region positions –1 and –3), suggesting a role in translational control. Mutations in this SMR were validated in whole-genome sequences of 7 cancer types, including 2 of 20 bladder cancers, 2 of 40 lung adenomas, and 3 of 172 breast cancers^3,19^. Bladder tumors with mutations in this SMR display altered RPS6KA1 (p90RSK) phosphorylation (*P* = 0.0005, *t-*test, Benjamini-Hochberg), a signal of increased cell-cycle proliferation^36^, and α -Tubulin (*P* = 4.3 × 10^−5^, *t-*test, Benjamini-Hochberg) levels, as determined by reverse-phase protein array (RPPA) assays^37^ (**Fig. 2f**, Supplementary Methods). These results establish the utility of WES data for identifying recurrently mutated non-coding regions and our SMR identification method in pinpointing potentially functional non-coding alterations in cancer.

### SMRs permit high-resolution analysis of protein coding alterations

As expected, most exome-derived SMRs lay within protein-coding regions, offering the opportunity to study recurrent somatic alterations within proteins. Although many protein domains share high burdens of somatic mutation in multiple cancers, protein domains can show remarkable cancer-type specific burdens of mutation as exemplified by VHL in kidney clear-cell carcinoma and SET in diffuse large B-cell lymphoma (**Fig. 3a**). The identification of SMRs across multiple cancer types permitted a systematic analysis of differential mutation frequencies with sub-genic and cancer specific resolution.

**Figure 3.**
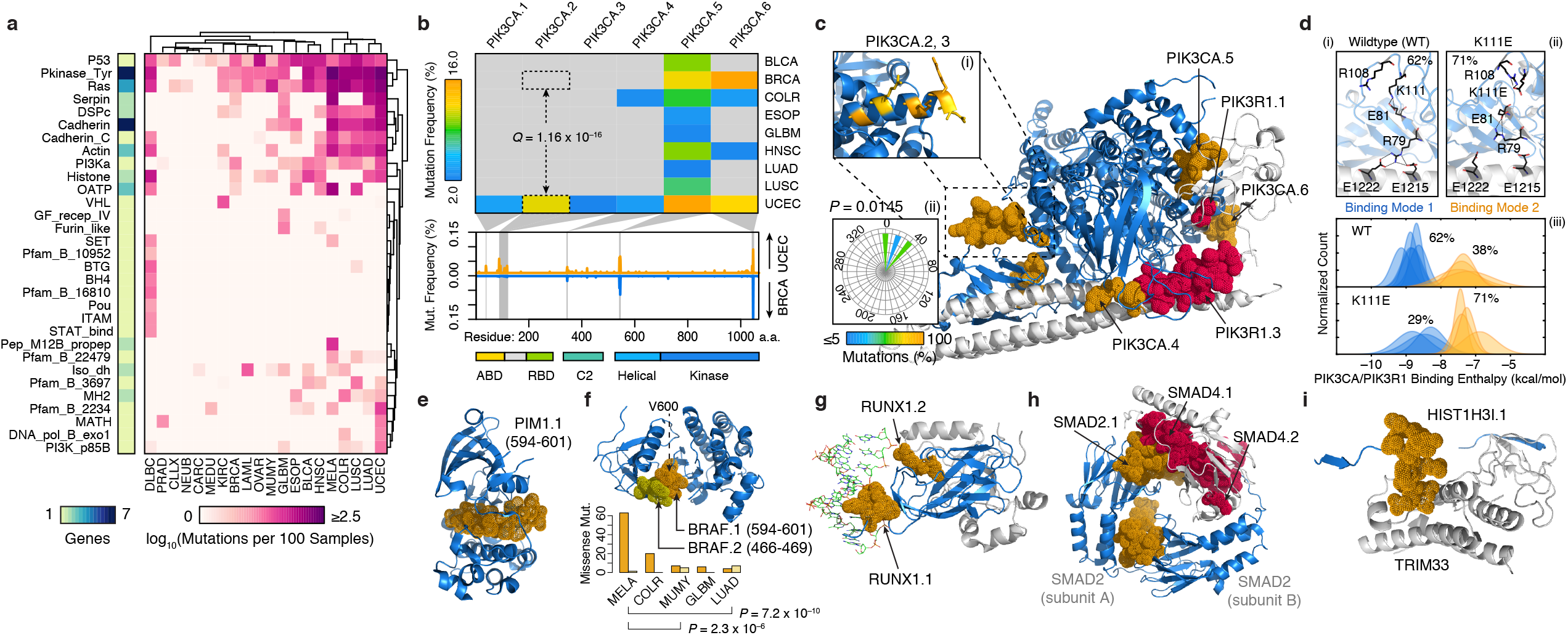
Structural mapping of SMRs onto proteins and complexes reveals regions differentially-altered among cancers and molecular interfaces targeted by recurrent alterations. (**a**) Matrix of non-synonymous mutations per PFAM protein domain, per cancer, per residue (right). Number of proteins per domain (left). (**b**) Mutation frequency matrix of PIK3CA SMRs across cancer types, and schematic comparison of per residue mutation frequency of PIK3CA domains^42^ in endometrial (UCEC; orange) and breast cancer (BRCA; blue) samples. Gray bars indicate SMRs within PIK3CA. (**c**) Co-crystal structure of the PIK3CA (p110α ; blue) and PIK3R1 (p85α ; gray) interaction (PDB: 2RDO, 2IUG, 3HIZ). Residues within endometrial cancer SMRs on PIK3CA (orange) and PIK3R1 (red) are rendered as solvent-accessible surfaces. Mutated residues within the PIK3CA.2, PIK3CA.3 SMR α-helix are colored yellow in (i), and their corresponding side-chain dihedral angles are shown (ii). (**d**) Large-scale simulations suggest PIK3CA–PIK3R1 binding is bimodal (iii). Mutations within the PIK3CA.2, PIK3CA.3 SMR α-helix interfere with R79 binding contacts at the PIK3R1 interface, as shown in wildtype (i), K111E (ii), and G118D (**Supplementary Fig. 8**). Molecular structures in an example of spatially-clustered mutations in diffuse large B-cell lymphoma (**e**; PIM1.1 (orange); PDB: 3CXW), spatially-clustered SMRs in multiple myeloma (**f**; BRAF.1 (orange) and BRAF.2 (yellow); PDB: 1UWH), a DNA (green) interface SMR on RUNX1 (**g**; RUNX1.1 (orange); PDB: 1H9D), reciprocal protein interface SMRs (**h**; SMAD2.1 (orange) and SMAD4.1, SMAD4.2 (red); PDB: 1U7V), and a histone H3.1 SMR (**i**; HIST1H3I.1 (orange); PDB:3U5N) in the TRIM33 interface. Structural alignments and molecular visualizations prepared with PyMOL (Schrödinger). The relative proportions of BRAF.1 and BRAF.2 missense mutations per cancer-type are shown in (**f**).

Among genes (*n*=94) with multiple SMRs, we detected 48 SMRs that are differentially mutated between cancer-types (**Supplementary Table 8**). A striking example of this differential targeting occurs within the catalytic subunit of the phosphoinositide 3-kinase, *PIK3CA* (*p110α*), a key oncogene implicated in a range of human cancers^38,39^. We detected six SMRs in *PIK3CA* across eight tumor-types (**Fig. 3b**), with multiple cancer types displaying SMRs in the helical (PIK3CA.5) and kinase (PIK3CA.6) domains. In contrast, we observed cancer-specific SMRs (PIK3CA.2, PIK3CA.3) affecting an α-helical region between the adaptor binding domain (ABD) and linker domains of PIK3CA. Up to 14% of uterine corpus endometrial carcinomas harbor alterations in these intron-separated SMRs although these regions are not highly recurrently altered in other cancers. For example, we observed significant (*Q* = 1.2 × 10^−16^, proportions test) differences in PIK3CA.2 alteration frequencies in endometrial and breast cancers (**Fig. 3b**), and further validated these differences (*P* = 0.02, proportions test) in whole-genome sequences^3,19^. These findings indicate that previously described differences^40^ in total PIK3CA mutation frequencies between endometrial and breast cancers could be localized to this region. Although the oncogenic effects of recurrent mutations in the ABD (PIK3CA.1), C2 (PIK3CA.4), helical (PIK3CA.5) and kinase (PIK3CA.5) domains of PIK3CA have been previously described^41–44^, mutations in this linker/ABD region remain unstudied. Interestingly, missense mutations within this region are directionally orientated (*P* = 0.0145, Rayleigh test) to one side of the α-helix, suggesting alterations to a molecular interface (**Fig. 3c**). Large-scale molecular dynamics simulations of PIK3CA–PIK3R1 indicate that PIK3CA.2 (K111E) and PIK3CA.3 (G118D) mutations can alter intermolecular salt bridge patterns at R79, which may result in a 1.8 kcal/mol loss of binding interactions compared to wildtype PIK3CA (**Fig. 3d**, **Supplementary Fig. 8**; Supplementary Methods). Taken together, these results suggest a previously unrecognized mechanism of oncogenic alteration in PIK3CA.

To systematically characterize the location of alterations with respect to three-dimensional protein structures, we leveraged structural information from 428 SMR-associated and known cancer-driver genes. We detected *n*=46 proteins with spatial (three-dimensional) clustering of missense mutations (**Supplementary Table 9**), as exemplified by PIM1, an SMR-associated serine/threonine kinase proto-oncogene (**Fig. 3e**; Supplementary Methods). This approach also identified spatial clustering between BRAF^*V*600^ and BRAF^*P-loop*^ SMRs (**Fig. 3f**), in which mutations have been shown to function through distinct mechanisms^45^. Moreover, we found that BRAF^*V*600^ mutations are more frequent in melanoma and colorectal cancers, whereas BRAF^*P-loop*^ mutations are more common in multiple myeloma and lung adenomas (*P* < 0.01, proportions test). In total, seven of 16 proteins with multiple SMRs displayed significant SMR spatial clustering (**Supplementary Table 10**), consistent with frequent spatial coherence in pathogenic alterations.

We next sought to identify SMRs that might affect the molecular interfaces of protein-protein and protein-DNA interactions, a recognized yet understudied mechanism of cancer-driver mutations^46–48^. We examined intermolecular distances between SMR residues and interacting proteins or DNA and identified 17 SMRs that likely alter molecular interfaces (**Table 1**; Supplementary Methods). These include 15 molecular interfaces of protein-protein and DNA-protein interactions with established cancer associations, such as the substrate-binding cleft of SPOP^49^, and DNA-binding interfaces on RUNX1 (**Fig. 3g**). We detected reciprocal SMRs at all electrostatic interfaces of the SMAD2–SMAD4 heterotrimer in colorectal cancer (**Fig. 3h**), as have been recently described^50^, and reciprocal SMRs at the regulatory PIK3CA–PIK3R1 interface in endometrial cancer (**Fig. 3b**). In addition, SMRs pinpoint recurrent alterations at the interface between histone H3.1 (**Fig. 3i**) and TRIM33, an E3 ubiquitin-protein ligase and transcriptional corepressor, and at the DNA-protein interface of histone H2B (**Supplementary Fig. 9**). These findings underscore and extend recent associations between altered epigenetic regulation and histone alterations in tumorigenesis^51^.

**Table 1.** Recurrently altered protein interfaces uncovered by SMRs.

| Protein (i) | Partner (j) | PDB | Chain (i) | Chain (j) | Region | Avg. Distance (Å) | Distance Ratio | Q-value | Status |
| --- | --- | --- | --- | --- | --- | --- | --- | --- | --- |
| VHL | TCEB1 | 3ZUN | I | H | chr3:10191469:10191513 | 7.259 | 0.395 | 7.62 x 10<sup>-10</sup> | Known |
| VHL | TCEB2 | 1LQB | C | A | chr3:10191469:10191513 | 9.867 | 0.367 | 7.62 x 10<sup>-10</sup> | Known |
| SPOP | H2AFY | 3HQH | A | M | chr17:47696421:47696467 | 7.962 | 0.462 | 3.72 x 10<sup>-8</sup> | Known |
| SMAD2 | SMAD4 | 1U7V | A | C | chr18:45374881:45374945 | 9.231 | 0.460 | 5.61 x 10<sup>-8</sup> | Known |
| HIST1H2BK | DNA | 2CV5 | D | J | chr6:27114446:27114519 | 9.730 | 0.520 | 3.27 x 10<sup>-7</sup> | Novel |
| TP53 | TP53BP1 | 1KZY | B | D | chr17:7578369:7578556 | 13.253 | 0.556 | 5.13 x 10<sup>-7</sup> | Known |
| SMAD4 | SMAD2 | 1U7V | B | C | chr18:48604665:48604797 | 11.878 | 0.694 | 5.13 x 10<sup>-7</sup> | Known |
| DNMT3A | DNMT3L | 2QRV | E | F | chr2:25463271:25463308 | 10.112 | 0.380 | 5.13 x 10<sup>-7</sup> | Known |
| SMAD4 | SMAD3 | 1U7F | B | C | chr18:48604665:48604797 | 11.883 | 0.700 | 1.94 x 10<sup>-6</sup> | Known |
| PIK3CA | PIK3R1 | 3HHM | A | B | chr3:178936070:178936099 | 9.028 | 0.335 | 2.56 x 10<sup>-6</sup> | Known |
| RUNX1 | DNA | 1H9D | C | H | chr21:36231782:36231792 | 8.957 | 0.351 | 0.001 | Known |
| HIST1H3I | TRIM33 | 3U5N | D | A | chr6:27839651:27840062 | 11.480 | 0.610 | 0.001 | Novel |
| HIST1H2BK | HIST1H4* | 2CV5 | D | F | chr6:27114446:27114519 | 13.680 | 0.664 | 0.002 | Novel |
| PPP2R1A | PPP2R5C | 2NPP | D | E | chr19:52716323:52716329 | 7.313 | 0.247 | 0.007 | Known |
| HRAS | RASA1 | 1WQ1 | R | G | chr11:534283:534291 | 5.302 | 0.350 | 0.007 | Known |
| PIK3R1 | PIK3CA | 3HIZ | B | A | chr5:67589138:67589149 | 6.713 | 0.567 | 0.008 | Known |
| NFE2L2 | KEAP1 | 2FLU | P | X | chr2:178098799:178098815 | 6.157 | 0.566 | 0.009 | Known |
| EGFR | EGF | 3NJP | B | A | chr7:55233035:55233043 | 8.763 | 0.386 | 0.019 | Known |
| FGFR2 | FGF8 | 2FDB | R | M | chr10:123279674:123279677 | 10.288 | 0.413 | 0.036 | Known |
| FBXW7 | SKP1 | 2OVR | B | C | chr4:153249384:153249385 | 9.352 | 0.346 | 0.036 | Known |
| FGFR2 | FGF2 | 1EV2 | H | A | chr10:123279674:123279677 | 11.685 | 0.406 | 0.037 | Known |

\*: Indicates multiple components partner proteins identified. "Status" indicates whether the SMR-harboring protein (i) is a known or novel cancer-driver gene.

### Molecular signature associations reveal the functional impact of SMR alterations

We sought to determine the potential functional impact of SMR alterations by their association with molecular signatures. Specifically, we leveraged RNA-seq, reverse-phase protein array (RPPA), and clinical data to ask whether: (1) SMRs alterations associate with distinct molecular signatures or survival outcomes, (2) SMR alterations correlate with similar molecular profiles in distinct cancers, same-gene SMR alterations associate with similar or different molecular signatures. These analyses provided mechanistic insights in how SMRs and the associated genes affect oncogenesis.

We found that mutations in SMRs were indeed associated with diverse changes in RNA expression, signaling pathways, and patient survival (**Fig. 4a**, **Supplementary Tables 11–14**; Supplementary Methods)^52^. These analyses revealed previously unappreciated connections between recurrent somatic mutations and molecular signatures. For example, synonymous point mutations in a bladder cancer SMR in sorting nexin 19 (*SNX19*; **Supplementary Table 12**) were associated with significant increases in protein expression levels of *RAB25* (*P* = 2.5 × 10^−27^, *t*-test; **Fig. 4b**), a RAS membrane trafficking GTPase that promotes ovarian and breast cancer progression, and is overexpressed in bladder cancer^53,54^. These increases are consistent with RNA expression differences of *RAB25* (*P* = 0.02; Wilcoxon rank sum test; **Fig. 4c**). Intriguingly, both SNX19 and RAB25 are implicated in intracellular trafficking, but the mechanism by which synonymous mutations in *SNX19* correlate with RAB25 expression remains to be determined.

**Figure 4.**
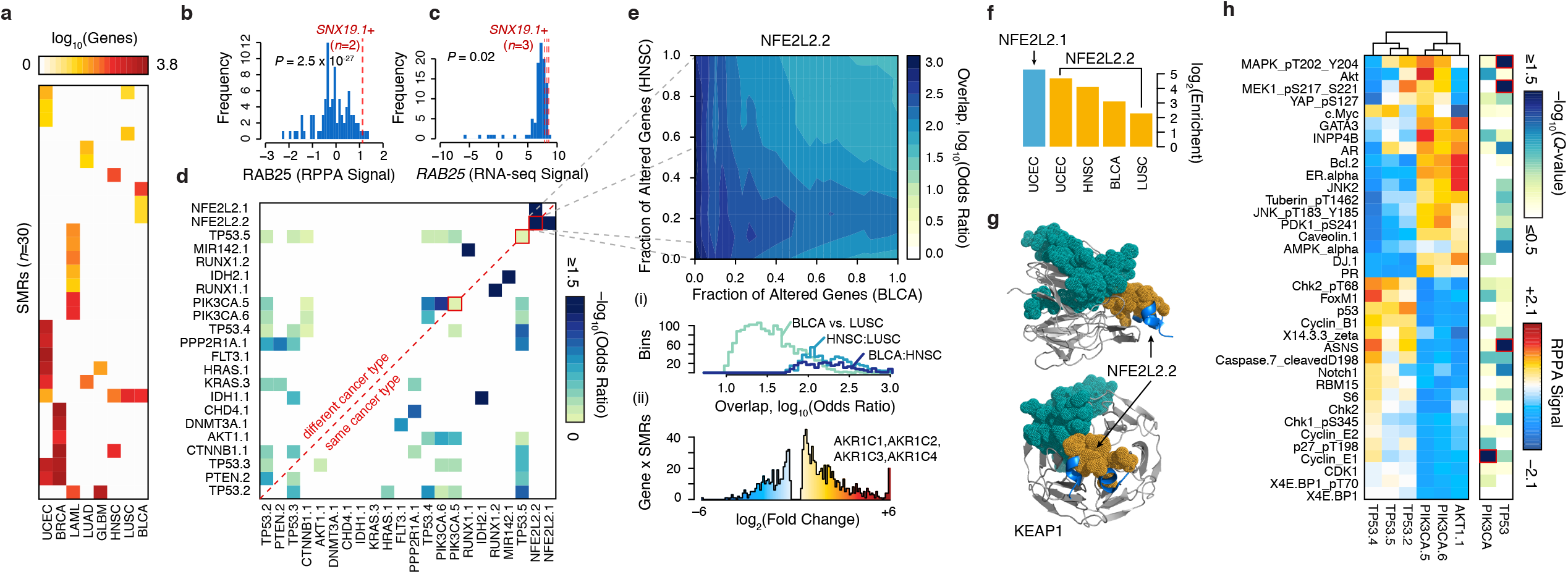
SMRs are associated with distinct molecular signatures. (**a**) Matched RNA-seq data for nine cancers revealed 30 distinct SMRs associated with at least 10 differentially expressed genes (FDR < 5%). Normalized reverse phase protein array (RPPA) (**b**) and RNA-seq (**c**) *RAB25* signals in *SNX19* SMR-altered (red) versus non-altered (blue) samples. Red lines indicate normalized signals for matched *SNX19* SMR-altered samples. (**d**) Similarity (Fisher’s exact test odds ratio; Supplementary Methods) between differentially expressed gene sets associated with mutations in each SMR pair. (**e**) Overlap of differentially expressed genes between patients with altered NFE2L2.2 in bladder cancer and head and neck carcinoma is shown. Differentially expressed genes were sorted by *p*-value and similarity was quantified by Fisher’s exact test odds ratio. (i) The distribution of odds ratios was summarized for three comparisons of gene expression profiles associated with NFE2L2.2 alterations across cancer types. (ii) Samples with NFE2L2.2 mutations exhibited highly increased expression of several aldo-keto reductase enzymes. (**f**) The relative enrichment for oxidoreductase activity (GO:0016616) among the differentially expressed genes in patients with NFE2L2 SMRs was plotted for specific cancer types (**Supplementary Table 13**). (**g**) Structure of SMR NFE2L2.2 (orange) in the KEAP1-binding domain (PDB: 3WN7). A sector of recurrent alterations on KEAP1 (teal) did not pass our 2% frequency cutoff. (**h**) Breast cancer patients were split into groups based on presence of mutations in six SMRs in PIK3CA, AKT1, and TP53. The median RPPA signal for these 36 markers is plotted along with the *q*-value (Kruskal-Wallis test) of differential expression between SMRs in TP53 or in PIK3CA. Markers with significant differential expression among intragenic SMRs were highlighted in red. Normalized RPPA-based expression was obtained from The Cancer Proteome Atlas (TCPA)^37^.

We identified concordant changes in gene expression between SMR pairs, revealing potential functional relationships among 23 SMRs from 17 genes (**Fig. 4d**). These included multiple well-established mechanistic relationships many of which were supported by RPPA measurements^37^, such as between *PIK3CA* and *AKT1*. Furthermore, this analysis revealed that mutations in the same SMR in different cancers can elicit similar molecular profiles in distinct cancers. For instance, we found that SMRs in the oncogenic transcription factor *NFE2L2^55^* were associated with large, concordant transcriptomic changes in four distinct cancer types (bladder, endometrial, lung squamous cell carcinoma, and head and neck cancer; **Fig. 4e**). The four genes with the highest increases in gene expression among endometrial cancer samples with alterations in *NFE2L2.1* were the aldo-keto reductases *AKR1C1-4* (**Fig. 4e**), which contribute to altered androgen metabolism and have been implicated in multiple cancer types^56–58^. Across all four cancer types, transcriptomic changes associated with *NFE2L2* SMR alterations were highly enriched for oxidoreductases acting on the CH-OH group of donors, NAD or NADP as acceptors (*P* ≤ 3.8 × 10^−2^, **Fig. 4f**). Mutations in KEAP1, a NFE2L2 binding partner, recapitulated the expression changes observed in patients with mutations in NFE2L2 SMRs (**Fig. 4g**; **Supplementary Fig. 10**; *P* < 0.01, Benjamini-Hochberg).

The identified SMRs also permitted interrogation of mutations in different regions of a given gene with respect to associated molecular signatures. For example in breast cancer, alterations in distinct SMRs within *PIK3CA* and *TP53* were associated with highly similar changes in protein-levels. Yet, we detected SMR-specific differences in cyclin E1 (CCNE1) levels among *PIK3CA* SMR-altered samples and ASNS levels and MAPK, MEK1 phosphorylation among *TP53* SMR-altered samples (**Fig. 4h**). These results establish intragenic differences in the molecular signatures of SMR alterations, and are consistent with pleiotropy in established oncogenes and tumor suppressors^59,60^.

### A large proportion of the structure in the distribution of cancer mutations remains unseen

SMR analysis leverages structure in the distribution of somatic driver mutations to identify cancer-associated coding and non-coding regions. We sought an alternative metric to assess the structure in the distribution of somatic coding mutations analyzed here by measuring the Gini coefficient of amino acid substitutions per residue in each cancer (**Fig. 5a**). Gini coefficients of dispersion were well-correlated with sample numbers (Spearman’s *ρ* = 0.74). Subsampling demonstrates that even with sample numbers in excess of 850, a large proportion of the structure of protein altering mutations –as measured by the Gini coefficient– in breast cancer remains unseen (**Fig. 5b**). These findings highlight the value of increasing tumor sample sizes in assessing the landscape of driver mutations.

**Figure 5.**
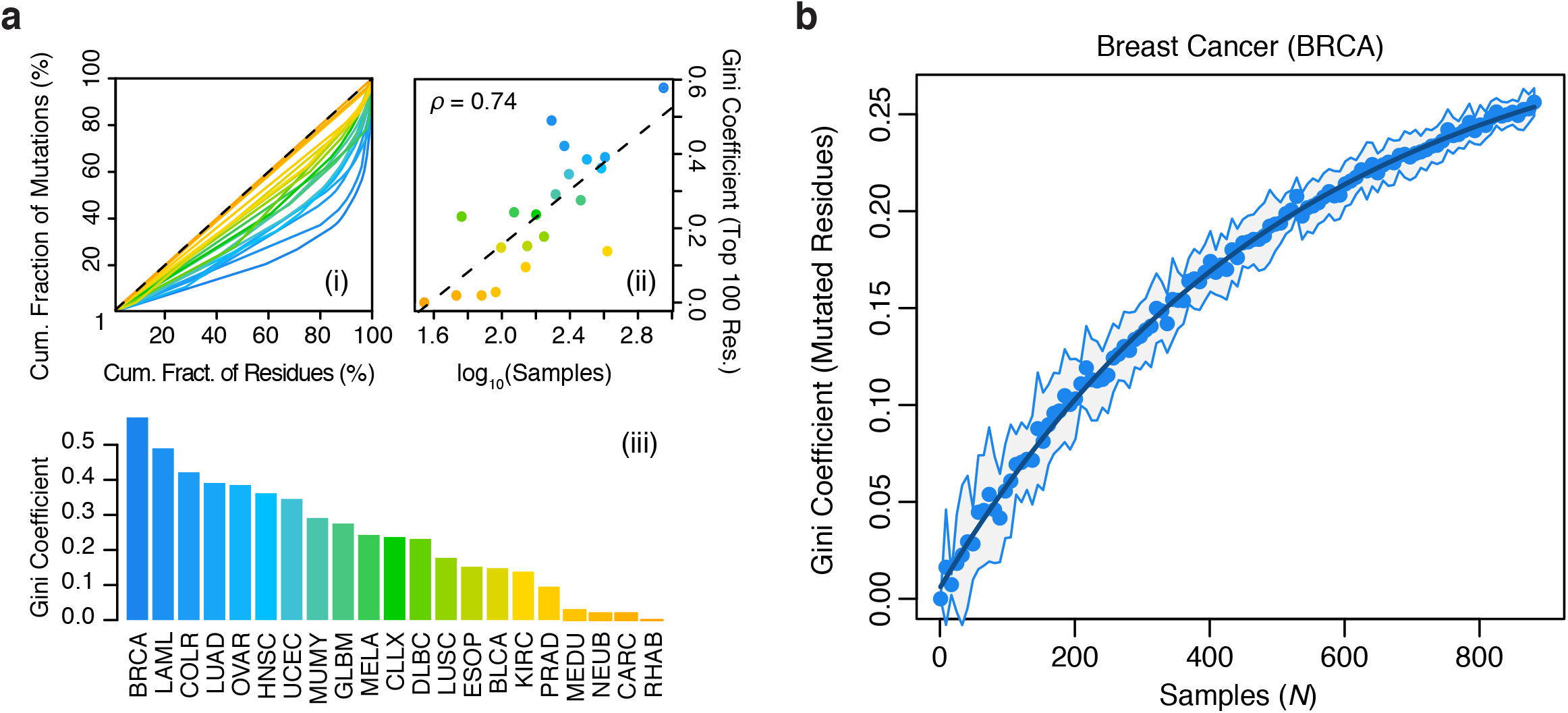
Structure in the distribution of cancer mutations remains largely uncharacterized. Gini coefficients of dispersion were calculated as the fraction of non-synonymous mutations contained per residue, across ∼19,000 proteins. (**a**) Lorenz curves (i), Gini-coefficients (iii), and their correlation with tumor sample numbers (ii) are shown. (**b**) Gini coefficients of non-synonymous mutation frequency in breast cancer as a function of (bootstrapped) sample size. Line of exponential fit is shown in dark blue. For comparisons between cancer types (**a**), the Gini coefficients were computed exclusively on the 100 most mutated residues per cancer.

## Discussion

With few exceptions, studies of disease-associated variation have focused on identifying predefined functional units with recurrent alterations in disease. This approach not only assumes accurate annotations but ignores the largely uncharacterized spectrum of functional elements that may be the targets of pathological variants. Our approach avoids these limitations and complements existing gene-level and pathway-based strategies for discovering cancer-drivers by identifying variably-sized, *significantly mutated regions* (SMRs) across 20 cancer types. SMR-associated genes include known cancer genes, such as *PIM1* and *MIR142* that were missed by gene-level analyses, as well as multiple novel genes with potential roles in cancer development.

Cancer SMRs target a diverse spectrum of functional elements in the genome, including single amino acids, complete coding exons and protein domains, miRNAs, 5’ UTRs, splice sites, and TF binding sites among others. Notably, several of the most frequently altered SMRs lay within non-coding regions. Mutation recurrence was validated in WGS data for non-coding SMRs at DHS sites in *KIAA0907* and *YAE1D1* promoters, and 5’ UTR alterations in *TBC1D12*, establishing the potential for discovering exon-proximal non-coding variation in WES. This functional diversity underscores both the varied mechanisms of oncogenic misregulation and the advantage of unbiased detection approaches.

Systematic analyses of SMRs within protein sequences and structures frequently provided mechanistic insight into altered molecular activities. We recovered many known cancer-implicated intermolecular interfaces, including recurrent alterations on opposing interfaces of PIK3CA– PIK3R1 and SMAD2–SMAD4. In addition, we observed NFE2L2 SMRs that reside in KEAP1 binding regions and result in concordant transcriptional changes across four distinct tumor types. Importantly, these transcriptional changes can be recapitulated by mutation of KEAP1, itself. We also uncovered recurrently altered histone interfaces, suggesting potential effects on global epigenetic dysregulation. For instance, histone H3.1 mutations at the TRIM33 interface may recapitulate TRIM33 loss-of-function and its associated pathogenic loss of SMAD4 transcriptional regulation^61^.

The detection of SMRs also permitted a sub-genic, cancer-specific analysis of somatic mutations and associated molecular signatures. Significant cancer-specific SMR mutation frequencies within BRAF, EGFR, and a functionally uncharacterized, directionally mutated α-helix in PIK3CA demonstrate substructure in the distribution of somatic mutations between cancers, a property which may arise from pleiotropic functions of macromolecules. The close geometric proximity and high directional uniformity, along with biophysical simulations, suggest that PIK3CA.2 and PIK3CA.3 mutations function through similar mechanisms. Taken together, the data implicates mutations in this α-helix in an elevated basal signaling activity of catalytic PIK3CA by way of weakened interactions with the regulatory PIK3R1 protein. Consistent with pleiotropic dependencies, alterations to SMRs within a single gene can be associated with distinct molecular signatures, as exemplified by both *PIK3CA* and *TP53* SMRs in breast cancers. Together, these results provide robust support for sub-genic functional targeting in distinct cancers and genes.

Yet, multiple possible methodological enhancements exist. For instance, SMR analysis would greatly benefit from refined measurements of background mutation frequencies, an area of active research^62–64^. Furthermore, three-dimensional protein folding and the interrupted nature of coding sequences (i.e. introns) present fundamental constraints on the efficacy of linear density clustering. This limitation, however, can be ameliorated through the generation of additional structural data across individual proteins and their conformational ensembles, allowing spatial (3D) clustering approaches to be applied to the large fraction of the proteome for which little structural information is currently available.

Although the sequencing of additional cancer genomes will further identification of novel cancer-driver genes^5^, characterizing the biochemical and cellular consequences of individual mutations is critical. We demonstrate that identifying the spatial concentration of mutations in the genome, when combined with additional genomic, biochemical, structural, or phenotypic information, often provides mechanistic insight into cancer etiology. The SMRs reported in this study provide a resource for the scientific community, nominating as functionally significant many novel elements. Applying recently-developed high-throughput approaches^65–67^ to directly interrogate sets of mutations found within SMRs may allow further understanding of the molecular mechanisms driving cancer and facilitate the development of more efficacious diagnostics and therapeutics. Our methods demonstrate valuable opportunities in complementing extant, gene-level approaches to identify pathogenic mutations with unbiased, multi-scale analysis of genomic variation. Finally, as the repertoire of functional elements in genomes continues to expand, SMRs provide a next-generation tool for increasingly large studies of genomic variation.

## URLs

Not applicable.

## Methods

Methods, additional display items, and their associated references are available in the online version of the article.

## Acknowledgements

We thank the TCGA, ICGC, and TCPA for making these large-scale cancer genomics and proteomic datasets available to the scientific community. C.A. was supported by NIH grants 3U54DK10255602 and 1P50HG00773501. C.C. was supported by the Child Health Research Institute, Lucile Packard Foundation for Children’s Health and the Stanford CTSA grant number UL1TR000093. J.A. was supported by NIH award 1U01HG007919-01. We thank H. Tang for discussions regarding statistical analyses. We thank M. M. Winslow, D. M. Fowler, S. Fields, and D. E. Webster for critical reading and suggestions to the manuscript.

### Author Contributions

C.L.A. and W.J.G. conceived of the project, and all authors designed experiments and methods. C.L.A. and C.C. developed methods for detection and analysis of significantly mutated regions (SMRs). C.L.A. constructed uniform annotations and non-Bayesian mutation probability models, and performed density-based clustering, scoring and empirical false discovery estimation (simulations), as well as regulatory (non-coding), structural (coding), frequency, and WGS recurrence analyses. C.C. constructed Bayesian mutation probability models, and performed RNA-seq, RPPA, and survival outcome analyses. G.K. carried out biophysical simulations, performed Hidden Markov Model based state decompositions, and computed binding enthalpies. C.L.A., C.C., J.A.R., G.K., M.P.S., and W.J.G. wrote the manuscript.

### Competing Financial Interests

M.P.S. is co-founder and a member of the scientific advisory board (SAB) of Personalis and a member of the SAB of Genapsys.

## References

1. Hodis, E. et al. A landscape of driver mutations in melanoma. Cell 150, 251–263 (2012).

2. Huang, F. W. et al. Highly recurrent TERT promoter mutations in human melanoma. Science 339, 957–959 (2013).

3. Alexandrov, L. B. et al. Signatures of mutational processes in human cancer. Nature (2013). doi:10.1038/nature12477

4. Lawrence, M. S. et al. Mutational heterogeneity in cancer and the search for new cancer-associated genes. Nature 499, 214–218 (2013).

5. Lawrence, M. S. et al. Discovery and saturation analysis of cancer genes across 21 tumour types. Nature 505, 495–501 (2014).

6. Ding, L., Wendl, M. C., McMichael, J. F. & Raphael, B. J. Expanding the computational toolbox for mining cancer genomes. Nat. Rev. Genet. 15, 556–570 (2014).

7. Davies, H. et al. Mutations of the BRAF gene in human cancer. Nature 417, 949–954 (2002).

8. Parsons, D. W. et al. An integrated genomic analysis of human glioblastoma multiforme. Science 321, 1807–1812 (2008).

9. Kane, D. P. & Shcherbakova, P. V. A common cancer-associated DNA polymerase ε mutation causes an exceptionally strong mutator phenotype, indicating fidelity defects distinct from loss of proofreading. Cancer Res. 74, 1895–1901 (2014).

10. Dees, N. D. et al. MuSiC: identifying mutational significance in cancer genomes. Genome Res. 22, 1589–1598 (2012).

11. Tamborero, D., Gonzalez-Perez, A. & Lopez-Bigas, N. OncodriveCLUST: exploiting the positional clustering of somatic mutations to identify cancer genes. Bioinformatics 29, 2238–2244 (2013).

12. Porta-Pardo, E. & Godzik, A. e-Driver: a novel method to identify protein regions driving cancer. Bioinformatics 30, 3109–3114 (2014).

13. Schnall-Levin, M., Zhao, Y., Perrimon, N. & Berger, B. Conserved microRNA targeting in Drosophila is as widespread in coding regions as in 3’UTRs. Proc. Natl. Acad. Sci. U. S. A. 107, 15751–15756 (2010).

14. Stergachis, A. B. et al. Exonic transcription factor binding directs codon choice and affects protein evolution. Science 342, 1367–1372 (2013).

15. Xiong, H. Y. et al. The human splicing code reveals new insights into the genetic determinants of disease. Science (2014). doi:10.1126/science.1254806

16. Wolfe, A. L. et al. RNA G-quadruplexes cause eIF4A-dependent oncogene translation in cancer. Nature 513, 65–70 (2014).

17. Gerstberger, S., Hafner, M. & Tuschl, T. A census of human RNA-binding proteins. Nat. Rev. Genet. (2014). doi:10.1038/nrg3813

18. ENCODE Project Consortium et al. An integrated encyclopedia of DNA elements in the human genome. Nature 489, 57–74 (2012).

19. Weinhold, N., Jacobsen, A., Schultz, N., Sander, C. & Lee, W. Genome-wide analysis of noncoding regulatory mutations in cancer. Nat. Genet. 46, 1160–1165 (2014).

20. Fredriksson, N. J., Ny, L., Nilsson, J. A. & Larsson, E. Systematic analysis of noncoding somatic mutations and gene expression alterations across 14 tumor types. Nat. Genet. (2014). doi:10.1038/ng.3141

21. Supek, F., Miñana, B., Valcárcel, J., Gabaldón, T. & Lehner, B. Synonymous mutations frequently act as driver mutations in human cancers. Cell 156, 1324–1335 (2014).

22. Leiserson, M. D. M. et al. Pan-cancer network analysis identifies combinations of rare somatic mutations across pathways and protein complexes. Nat. Genet. (2014). doi:10.1038/ng.3168

23. Araya, C. L. et al. Regulatory analysis of the C. elegans genome with spatiotemporal resolution. Nature 512, 400–405 (2014).

24. Stergachis, A. B. et al. Conservation of trans-acting circuitry during mammalian regulatory evolution. Nature 515, 365–370 (2014).

25. Roadmap Epigenomics Consortium et al. Integrative analysis of 111 reference human epigenomes. Nature 518, 317–330 (2015).

26. Cingolani, P. et al. A program for annotating and predicting the effects of single nucleotide polymorphisms, SnpEff: SNPs in the genome of Drosophila melanogaster strain w1118; iso-2; iso-3. Fly 6, 80–92 (2012).

27. Martin Ester, Hans-peter Kriegel, Jörg S, Xiaowei Xu. A density-based algorithm for discovering clusters in large spatial databases with noise. KDD (1996).

28. Cenik, C. et al. Genome analysis reveals interplay between 5’UTR introns and nuclear mRNA export for secretory and mitochondrial genes. PLoS Genet. 7, e1001366 (2011).

29. Andrew Futreal P., et al. A census of human cancer genes. Nat. Rev. Cancer 4, 177–183 (2004).

30. Santarius, T., Shipley, J., Brewer, D., Stratton, M. R. & Cooper, C. S. A census of amplified and overexpressed human cancer genes. Nat. Rev. Cancer 10, 59–64 (2010).

31. Malhotra, A. et al. Breakpoint profiling of 64 cancer genomes reveals numerous complex rearrangements spawned by homology-independent mechanisms. Genome Res. 23, 762–776 (2013).

32. Jäger, D. et al. Identification of a tissue-specific putative transcription factor in breast tissue by serological screening of a breast cancer library. Cancer Res. 61, 2055–2061 (2001).

33. Mei, Y.-P. et al. Small nucleolar RNA 42 acts as an oncogene in lung tumorigenesis. Oncogene 31, 2794–2804 (2012).

34. Budinska, E. et al. Gene expression patterns unveil a new level of molecular heterogeneity in colorectal cancer. J. Pathol. 231, 63–76 (2013).

35. Uhlén, M. et al. Proteomics. Tissue-based map of the human proteome. Science 347, 1260419 (2015).

36. Lara, R., Seckl, M. J. & Pardo, O. E. The p90 RSK family members: common functions and isoform specificity. Cancer Res. 73, 5301–5308 (2013).

37. Li, J. et al. TCPA: a resource for cancer functional proteomics data. Nat. Methods 10, 1046–1047 (2013).

38. Samuels, Y. et al. High frequency of mutations of the PIK3CA gene in human cancers. Science 304, 554 (2004).

39. Thorpe, L. M., Yuzugullu, H. & Zhao, J. J. PI3K in cancer: divergent roles of isoforms, modes of activation and therapeutic targeting. Nat. Rev. Cancer 15, 7–24 (2014).

40. Cancer Genome Atlas Research Network et al. Integrated genomic characterization of endometrial carcinoma. Nature 497, 67–73 (2013).

41. Miled, N. et al. Mechanism of two classes of cancer mutations in the phosphoinositide 3-kinase catalytic subunit. Science 317, 239–242 (2007).

42. Huang, C.-H. et al. The structure of a human p110alpha/p85alpha complex elucidates the effects of oncogenic PI3Kalpha mutations. Science 318, 1744–1748 (2007).

43. Huang, C.-H., Mandelker, D., Gabelli, S. B. & Amzel, L. M. Insights into the oncogenic effects of PIK3CA mutations from the structure of p110alpha/p85alpha. Cell Cycle 7, 1151–1156 (2008).

44. Gkeka, P. et al. Investigating the Structure and Dynamics of the PIK3CA Wild-Type and H1047R Oncogenic Mutant. PLoS Comput. Biol. 10, e1003895 (2014).

45. Haling, J. R. et al. Structure of the BRAF-MEK complex reveals a kinase activity independent role for BRAF in MAPK signaling. Cancer Cell 26, 402–413 (2014).

46. Kar, G., Gursoy, A. & Keskin, O. Human cancer protein-protein interaction network: a structural perspective. PLoS Comput. Biol. 5, e1000601 (2009).

47. Ghersi, D. & Singh, M. Interaction-based discovery of functionally important genes in cancers. Nucleic Acids Res. 42, e18 (2014).

48. Cheng, F. et al. Studying tumorigenesis through network evolution and somatic mutational perturbations in the cancer interactome. Mol. Biol. Evol. 31, 2156–2169 (2014).

49. Barbieri, C. E. et al. Exome sequencing identifies recurrent SPOP, FOXA1 and MED12 mutations in prostate cancer. Nat. Genet. 44, 685–689 (2012).

50. Fleming, N. I. et al. SMAD2, SMAD3 and SMAD4 mutations in colorectal cancer. Cancer Res. 73, 725–735 (2013).

51. Yuen, B. T. K. & Knoepfler, P. S. Histone H3.3 mutations: a variant path to cancer. Cancer Cell 24, 567–574 (2013).

52. Hornbeck, P. V. et al. PhosphoSitePlus: a comprehensive resource for investigating the structure and function of experimentally determined post-translational modifications in man and mouse. Nucleic Acids Res. 40, D261–70 (2012).

53. Cheng, K. W. et al. The RAB25 small GTPase determines aggressiveness of ovarian and breast cancers. Nat. Med. 10, 1251–1256 (2004).

54. Zhang, J. et al. Overexpression of Rab25 contributes to metastasis of bladder cancer through induction of epithelial-mesenchymal transition and activation of Akt/GSK-3β/Snail signaling. Carcinogenesis 34, 2401–2408 (2013).

55. DeNicola, G. M. et al. Oncogene-induced Nrf2 transcription promotes ROS detoxification and tumorigenesis. Nature 475, 106–109 (2011).

56. Ji, Q. et al. Selective loss of AKR1C1 and AKR1C2 in breast cancer and their potential effect on progesterone signaling. Cancer Res. 64, 7610–7617 (2004).

57. Stanbrough, M. et al. Increased expression of genes converting adrenal androgens to testosterone in androgen-independent prostate cancer. Cancer Res. 66, 2815–2825 (2006).

58. Rižner, T. L., Šmuc, T., Rupreht, R., Šinkovec, J. & Penning, T. M. AKR1C1 and AKR1C3 may determine progesterone and estrogen ratios in endometrial cancer. Mol. Cell. Endocrinol. 248, 126–135 (2006).

59. Zhao, L. & Vogt, P. K. Helical domain and kinase domain mutations in p110a of phosphatidylinositol 3-kinase induce gain of function by different mechanisms. Proceedings of the National Academy of Sciences 105, 2652–2657 (2008).

60. Wu, X. et al. Activation of diverse signalling pathways by oncogenic PIK3CA mutations. Nat. Commun. 5, 4961 (2014).

61. Hatakeyama, S. TRIM proteins and cancer. Nat. Rev. Cancer 11, 792–804 (2011).

62. Polak, P. et al. Cell-of-origin chromatin organization shapes the mutational landscape of cancer. Nature 518, 360–364 (2015).

63. Supek, F. & Lehner, B. Differential DNA mismatch repair underlies mutation rate variation across the human genome. Nature (2015). doi:10.1038/nature14173

64. Reijns, M. A. M. et al. Lagging-strand replication shapes the mutational landscape of the genome. Nature (2015). doi:10.1038/nature14183

65. Fowler, D. M. et al. High-resolution mapping of protein sequence-function relationships. Nat. Methods 7, 741–746 (2010).

66. Buenrostro, J. D. et al. Quantitative analysis of RNA-protein interactions on a massively parallel array reveals biophysical and evolutionary landscapes. Nat. Biotechnol. (2014). doi:10.1038/nbt.2880

67. Guenther, U.-P. et al. Hidden specificity in an apparently nonspecific RNA-binding protein. Nature (2013). doi:10.1038/nature12543

